# Comparison of extracellular vesicles isolation methods reveals method-dependent protein and miRNA profiles in saliva

**DOI:** 10.1101/2025.11.26.690666

**Authors:** Elodie Simphor, Jérémy Boulestreau, Romane Marchal, Franck Molina, Malik Kahli

## Abstract

Salivary extracellular vesicles (EVs) represent a powerful, non-invasive source of biomarkers for disease diagnosis and monitoring. Their molecular cargo reflects systemic and local physiological states, offering a window into neurological and inflammatory disorders. However, the diversity of EV isolation protocols and the possibility that each enriches distinct EV subpopulations remains a major barrier to reproducibility and data comparability. We conducted a comprehensive comparison of three EV isolation methods: ultracentrifugation (UC), PEG-based precipitation (Q), and immunoaffinity capture (M) to evaluate their impact on EV yield, purity, and molecular composition. Salivary EVs from healthy volunteers were analysed using proteomic and small-RNA sequencing approaches. Principal component analysis revealed clear isolation method-dependent clustering, where M-derived EVs displayed the most distinct profile. UC and Q produced broader proteomic repertoires with higher total protein content, whereas M-isolated EVs exhibited greater purity and enrichment of trafficking-and lysosome-associated proteins. Over 731 miRNAs selected, 28 were consistently altered across methods and 65 uniquely enriched in M isolates. RT-qPCR confirmed key directional trends. These 93 method-dependent miRNAs have predicted targets associated with synaptic structure and neurodegenerative pathways. These findings show that isolation methodology deeply shapes salivary EV cargo and suggest that immunoaffinity capture can isolate specific EV populations meeting diagnostic requirements.

## Introduction

Extracellular vesicles (EVs) are membrane-bound particles released by most cell types, playing key roles in intercellular communication ^1^. EVs include exosomes, microvesicles (ectosome), and apoptotic bodies, which differ in size and biogenesis but share the capacity to transfer molecular cargo (such as proteins, lipids, metabolites and RNA such as microRNAs (miRNAs)) to recipient cells ^2^. This exchange influences physiological processes, including immune response ^3^, cell proliferation ^4^, and tissue repair^5^, and contributes to pathological mechanisms like cancer progression ^6^ and neurodegeneration ^7^.

The biomedical potential of EVs has attracted growing interest, particularly for their use as biomarkers. EV cargo reflects the physiological state of their cells of origin ^2^, positioning them as candidates for so-called “liquid biopsies”: non-invasive assays that can provide diagnostic, prognostic, and monitoring information ^8,9^. EVs have been successfully isolated from blood ^10^, urine ^11^, cerebrospinal fluid ^12^, and saliva ^13^. Among these, saliva is especially promising due to its ease of collection, non-invasive nature, and reduced risk of blood contamination. Salivary EVs may provide insights into both local and systemic health conditions, such as oral cancer ^14^, Sjögren’s syndrome ^15,16^, and neurodegenerative disorders like Alzheimer’s ^17^ or Parkinson’s ^18,19^. Proteomic and transcriptomic changes in salivary EVs have already been linked to early pathological alterations, underscoring their diagnostic potential ^16,19,20^.

Despite these advantages, isolating EVs from saliva presents unique challenges. Saliva contains abundant proteins, mucins, enzymes, bacterial vesicles, and other contaminants that complicate isolation and may compromise EV purity ^21^. Moreover, EV populations are heterogeneous, varying in size and cargo composition, which complicates reproducibility across studies.

Several methods are commonly employed to isolate EVs, each offering distinct advantages and limitations. Ultracentrifugation (UC), historically regarded as the gold standard, separates vesicles based on size and density but is labor-intensive, requires specialized equipment, and may co-isolate protein aggregates and non-vesicular particles ^22,23^. Precipitation approaches, such as polyethylene glycol (PEG)-based co-precipitation, are scalable and user-friendly but often capture soluble proteins and lipoproteins, compromising downstream analyses ^24–26^. Immuno-affinity capture, which targets tetraspanins such as CD9, CD63, and CD81, provides high specificity but yields fewer vesicles and may restrict recovery to selected EV subpopulations ^27^. Thus, each method involves trade-offs between yield, purity, and scalability, and the optimal approach depends on the intended downstream application; whether the goal is molecular cargo characterization (e.g., proteomic, transcriptomic, or lipidomic profiling), evaluation of EV functionality or biodistribution, or biomarker discovery in clinical contexts.

Proteomic and miRNA analyses are central to understanding EV composition and function, as well as their potential as biomarkers. Proteomics enables the identification of EV-associated proteins such as membrane receptors, signalling molecules, and cytoskeletal components that reflect cellular origin and biological activity. However, proteomic profiles can be influenced by co-isolated non-vesicular proteins, making sample purity a key determinant of interpretability ^26^. Similarly, EV-carried miRNAs are remarkably stable and clinically informative ^28–30^, but accurate quantification depends on efficient separation of vesicular and non-vesicular RNA fractions. Although both approaches provide complementary insights into EV biology, how different isolation methods shape these molecular readouts (particularly in saliva) remains insufficiently characterized. Understanding these effects is essential to improve reproducibility and ensure that biomarker signals accurately reflect vesicle biology rather than technical bias ^31–33^.

Although systematic evaluations of EV isolation have been performed in plasma ^33–35^, urine ^36–38^ and CSF ^39,40^, comparable analyses for saliva remain scarce. Recent work has begun to highlight the sensitivity of salivary EV profiles to isolation method ^26,41,42^, but comprehensive multi-omics comparisons remain limited. Given saliva’s diagnostic promise and complexity, establishing best practices for EV isolation is essential.

In a previous study, we compared UC, PEG-based precipitation, and immunoaffinity capture for salivary EVs, assessing yield, purity, protein markers, and selected miRNAs, and also examined the effect of filtration ^43^. The present study expands the analysis to include comprehensive proteomic and small RNA sequencing of EVs obtained by these methods. By integrating proteomic and transcriptomic readouts, we assess how isolation strategy shapes the salivary EV molecular landscape. Our goal is to provide practical data that inform method selection, support standardization, and facilitate reproducibility in biomarker discovery and clinical translation.

## Methods

### Saliva collection and sample processing

Unstimulated whole saliva was collected in the morning (9:00–10:00 AM) from healthy male volunteers (18–40 years old) who had refrained from eating, drinking, or smoking for at least 1 hour. Participants passively drooled into sterile tubes, and a minimum of 4 mL was collected per individual. Samples showing visible blood contamination were excluded. Saliva was placed on ice and processed within 1 hour: first centrifuged at 300 × g for 10 min at 4 °C to remove cells, followed by 3000 × g for 30 min at 4 °C to eliminate residual debris. The resulting supernatant was used for EV isolation. All samples were processed individually (not pooled), and each “n” corresponds to a unique biological replicate. Ethical approval was granted by a French national committee (CPP-NORD OUEST III) on April 21, 2023 (23.00930.000169), and all participants provided written informed consent. The study was registered at www.clinicaltrials.gov (NCT06149351).

### EV isolation and concentration

Three methods were used to isolate EVs from 1 mL of whole saliva supernatant (WS): ultracentrifugation, co-precipitation, and immunoaffinity capture.

**Ultracentrifugation (UC):** WS was diluted in 24 mL of PBS and ultracentrifuged at 100,000 × g for 1 h at 4 °C (JXN-30, JA-30.50 Ti rotor, Beckman Coulter). The pellet was resuspended in 10 mL PBS and centrifuged again under the same conditions. The final pellet was resuspended in 100 µL PBS and either analysed immediately or stored at –80 °C.

**Co-precipitation (Q):** EVs were isolated using the miRCURY® Exosome Kit (Qiagen, Cat. #76743). Briefly, 400 µL of precipitation buffer was added to 1 mL of WS, incubated at 4 °C for 1 h, and centrifuged at 10,000 × g for 30 min at room temperature. The pellet was resuspended in 100 µL of the provided buffer and stored on ice or at –80 °C.

**Immunoaffinity capture (M):** EVs were isolated using the Exosome Isolation Kit Pan, Human (Miltenyi Biotec, Cat. #130-111-572). 50 µL of MicroBeads (anti-CD9, CD63, CD81) were incubated with 1 mL of WS for 1 h at room temperature with agitation. The mixture was loaded onto a µMACS column, washed, and EVs were eluted in 100 µL of buffer. Bead-only controls were processed in parallel to assess background. Eluted EVs were used fresh or stored at –80 °C.

### EV characterization

EVs were characterized according to ISEV (MISEV2023) guidelines ^44^. Size distribution and concentration were assessed using Nanoparticle Tracking Analysis (NTA; NanoSight NS300, Malvern Panalytical) in PBS. Measurements (n ≥ 3) were acquired with NTA software v3.4 (camera level 16; detection threshold 5), and analysed to determine mean size and particle counts.

Cryo-transmission electron microscopy (cryo-TEM) was performed by depositing EVs on glow-discharged Lacey grids, blotting, and vitrifying in liquid ethane using a CP3 cryo-plunge system (Gatan). Grids were imaged on a JEOL 2200FS FEG TEM (200 kV, 4k × 4k CCD, 20 eV energy filter) under low-dose conditions.

Protein concentration was measured using the Micro BCA Protein Assay Kit (ThermoFisher Scientific) at 562 nm (Synergy H1, BioTek) and quantified against BSA standards (µg/µL). Western blotting for EV marker profiling is described in the next section. All EV samples were used fresh or stored at −80 °C to preserve integrity.

### SDS-PAGE and Western blot

EV samples were lysed in RIPA buffer with protease inhibitors (Roche) and Laemmli buffer containing 2.5% β-mercaptoethanol. Equal amounts of protein (20 µg) were loaded on 12.5% SDS-polyacrylamide gels and separated at 200 V for 50 min. Proteins were transferred to nitrocellulose membranes (Trans-Blot, Bio-Rad) at 90 V for 1 h. Membranes were blocked with 5% skim milk in PBS-Tween 20 (0.1%) for 1 h and incubated overnight at 4°C with primary antibodies (Abcam): CD9 (ab263019), CD63 (ab134045), HSP70 (ab181606), TSG101 (ab125011) at 1:1000, and albumin (ab19180) at 1:20,000.

After washing, membranes were incubated with HRP-conjugated secondary antibodies (Sigma) for 1 h at room temperature. Signal detection was performed using Clarity Max ECL substrate (Bio-Rad), and imaging was done with a ChemiDoc MP system (Bio-Rad). Band intensities were quantified with ImageJ.

### Proteomics

Proteomic analysis was performed at Plateforme de Proteomique Fonctionnelle de Montpellier (FPP on triplicates of WS, EV UC, EV Q, and EV M samples (14.5 µg protein per sample). Proteins were digested using S-Trap™ micro spin columns (Protifi) following the manufacturer’s protocol with minor modifications. Briefly, samples were solubilized in 5% SDS, reduced with 20 mM DTT (10 min, 95°C), alkylated with 40 mM IAA (30 min, dark), acidified with 1.2% phosphoric acid, and diluted in binding buffer (100 mM TEABC in 90% methanol). Proteins were trapped by centrifugation (4000 g, 1 min), washed five times, and digested with 1 µg trypsin (Promega) for 2 h at 47°C. Peptides were eluted, vacuum-dried, and stored for MS analysis.

Peptide separation and analysis were performed via nanoLC-MS/MS using a Q-Exactive HF mass spectrometer (ThermoFisher) coupled with an Ultimate 3000 RSLC system. After online desalting on a PepMap® precolumn, peptides were separated on a PepMap® C18 column using a 123-min gradient (2–40% acetonitrile, 0.1% formic acid, 300 nL/min). MS1 scans were acquired in the Orbitrap (375–1500 m/z, resolution 60,000), followed by MS2 (Top12, HCD fragmentation, resolution 30,000).

Raw spectra were processed with MaxQuant v2.0.3.0 ^45^ using the Andromeda search engine, label-free quantification (LFQ), and iBAQ enabled ^46^. Spectra were searched against the UniProt *Homo sapiens* reference proteome (release 2023_03) with common contaminants and decoy sequences included. Fixed and variable modifications included carbamidomethylation (Cys), oxidation (Met), and N-terminal acetylation. Peptide and protein FDRs were set at 1%. Data visualization and statistical analysis were performed using Perseus v1.6.15.0.^47^.

### RNA extraction, RT-qPCR, and small RNA sequencing

Total small RNA was extracted from EV and WS samples using the miRNeasy Serum/Plasma Kit (Qiagen, #217184) and eluted in 14 µL of RNase-free water. RNA yield was quantified using the Qubit™ microRNA Assay Kit (ThermoFisher, #Q32881), and size distribution was assessed via the LabChip GX system (Caliper-PerkinElmer) using the Small RNA Assay (#CLS153530) to confirm the presence of miRNAs.

For RT-qPCR, 2 ng of RNA was reverse-transcribed using the miRCURY® LNA® RT Kit (Qiagen). qPCR was conducted on 20 pg of cDNA using miRNA-specific primers (Supplementary Table S1) and the miRCURY® LNA® SYBR® Green PCR Kit (Qiagen).

Small RNA libraries were prepared from 10 ng of miRNA using the TruSeq Small RNA Library Prep Kit (Illumina, #RS-200-0012). After adapter ligation, cDNA synthesis, and PCR amplification (11 cycles), libraries were purified with Ampure XP beads (Beckman Coulter, #A63880), assessed on the LabChip GX system, and quantified via Qubit™ dsDNA HS Assay (#Q32851). Equimolar pooling was followed by sequencing on the NextSeq 500 platform using the high output v2.5 kit (1.8 pM, #20024906).

Post-sequencing, quality control (FastQC, multiQC), adapter trimming, and Phred score filtering were applied. Reads were mapped to the human genome (GRCh38.p14) and miRBase v20 using the miRDeep2 pipeline, enabling both miRNA identification and quantification.

### Bioinformatics and Statistical Analysis

#### Proteomics

Label-free quantification (LFQ) data from MaxQuant (v2.0.3.0) were used to analyse protein abundance across WS, UC-, Q-, and M-derived EVs. Data were filtered to exclude contaminants, reverse hits, single-peptide identifications, and inconsistently quantified proteins. LFQ intensities were log₂-transformed and missing values were imputed using the “missing not at random” (MNAR) method (Gaussian distribution, 1.8 SD shift, width 0.3). Differential expression was assessed using the limma package (R Bioconductor), applying empirical Bayes statistics. Significance was defined as |log₂FC| ≥ 1 with FDR-adjusted p < 0.05. Venn diagrams were generated using an online tool (https://bioinformatics.psb.ugent.be/webtools/Venn/).

#### miRNA-seq

Raw sequencing data were processed with FastQC (v0.12.1) and multiQC (v1.23). Adapter trimming and quality filtering (Phred score > 20) were performed using fastp (v0.23.4). Reads were mapped to GRCh38.p14 and annotated with miRBase v20 using the miRDeep2 pipeline (v0.13). miRNA expression quantification and differential analysis were performed with DESeq2 (v1.42.1) ^48^, using a Wald test under a negative binomial GLM. Multiple testing correction was done using the Benjamini-Hochberg method. Normalized counts (CPM) were calculated via edgeR (v4.0.16).

#### Functional Enrichment

Target gene prediction and validation were conducted using the multiMiR R package (v1.24.0). Gene ontology (GO) and KEGG pathway enrichment analyses were performed using clusterProfiler (v4.10.1).

#### General Statistics

All other statistical analyses were performed in GraphPad Prism v10. Data were tested for normality. Wilcoxon signed-rank, Friedman’s, or Kruskal-Wallis tests were applied as appropriate. Dunn’s post hoc test was used for multiple comparisons. Results are reported as mean ± SEM. Significance was set at *p* < 0.05.

## Results

### Experimental workflow and EV characterization

As described previously ^43^, unstimulated whole saliva was collected from healthy male volunteers and processed immediately (Fig. 1). After sequential low-speed centrifugation, 500 µL of clarified saliva (whole saliva, WS) was retained for direct analysis, while 1 mL aliquots were subjected to EV isolation by ultracentrifugation (UC), PEG-based precipitation (Q), or immunoaffinity capture (M). EV preparations were either analysed fresh or stored at −80 °C. Characterization included nanoparticle tracking analysis (NTA), cryo-electron microscopy (cryo-EM), protein quantification, and Western blotting (WB).

**Figure 1:**
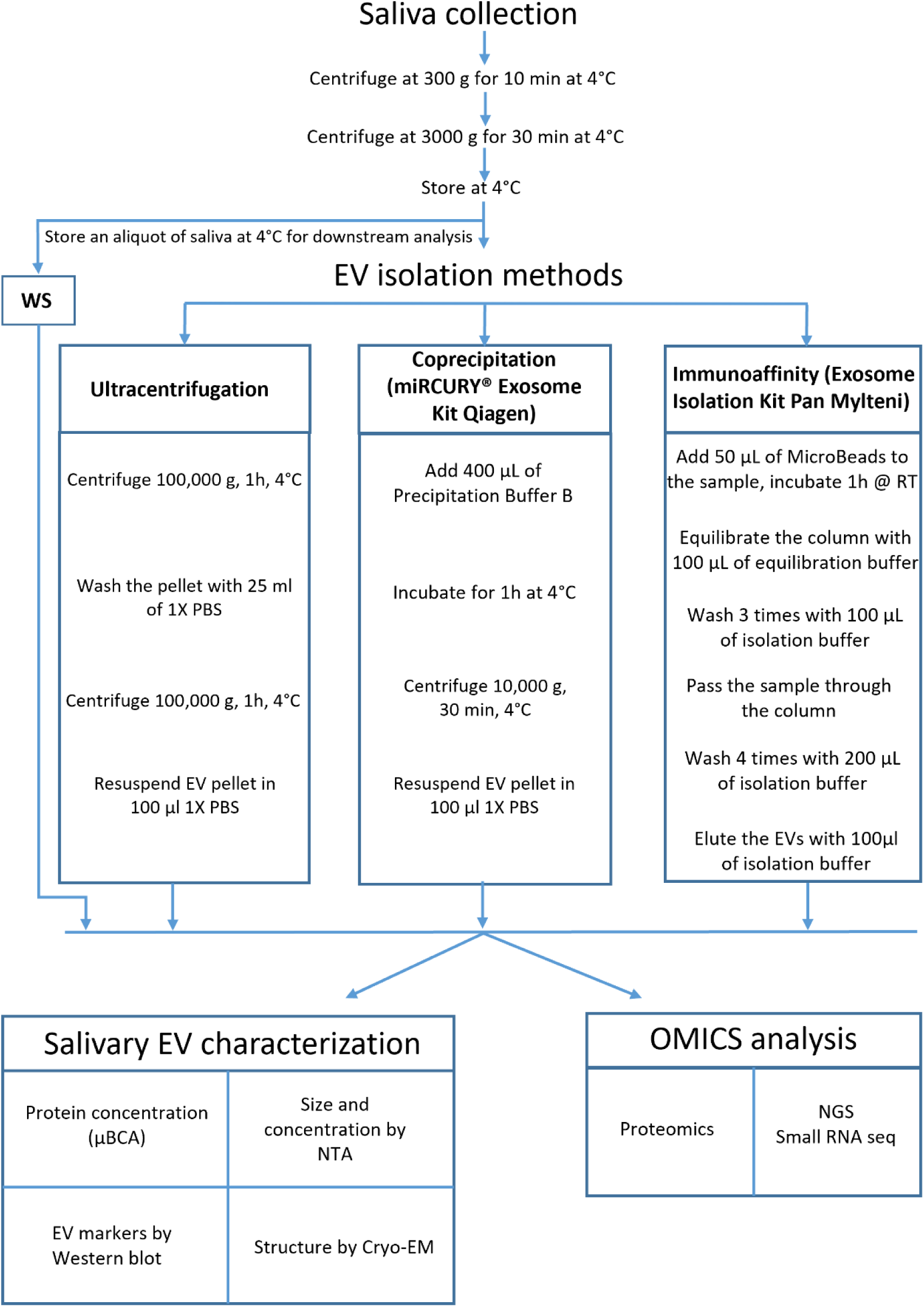
Schematic overview of the experimental workflow for salivary EV isolation and analysis. Unstimulated saliva (4 mL) was collected from healthy donors and sequentially centrifuged at 300 × *g* and 3000 × *g* to remove cells, large debris, and residual contaminants. The resulting whole saliva supernatant (WS) was divided into four aliquots: 0.5 mL was retained as the WS control, and 1 mL was allocated to each of the three EV isolation methods: ultracentrifugation (UC), polymer-based precipitation (Q), and immunoaffinity capture (M). Isolated EVs were subsequently characterized and subjected to proteomic and small RNA sequencing analyses.

NTA showed that UC and Q yielded EVs of similar size distributions, although Q isolates had a smaller mode diameter (147 nm vs. 180 nm) and lower particle concentration (1.9 × 10⁸ vs. 4.5 × 10⁸ particles/mL). Protein quantity was higher in Q samples, resulting in a reduced particle-to-protein ratio. M-isolated EVs displayed the highest particle counts (1.5 × 10⁹ particles/mL) and smallest diameters (mean 84 nm; mode 62 nm), with a purity index close to that of UC (Table 1, Fig. 2A, 2C, 2D).

**Figure 2:**
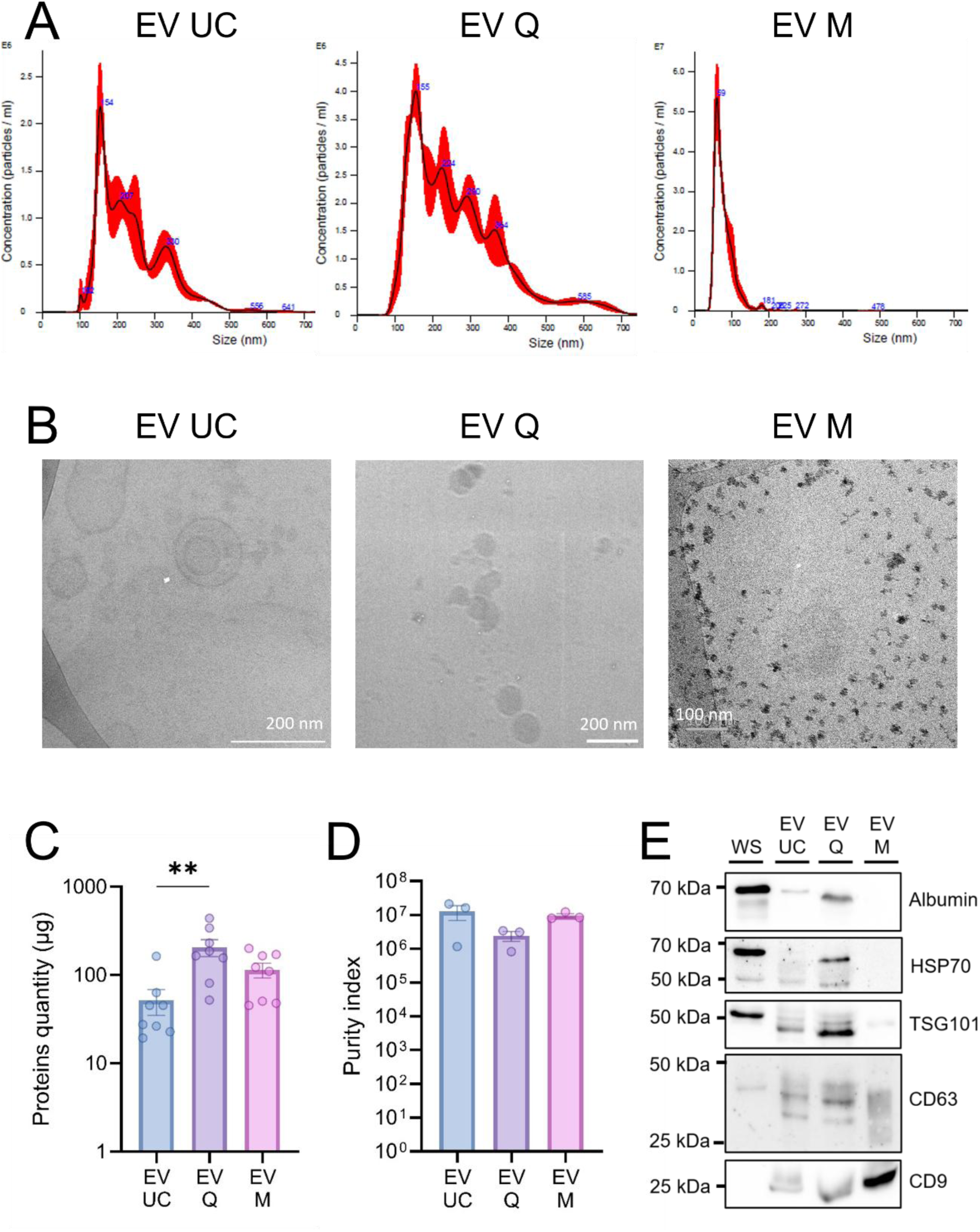
Characterization of human salivary extracellular vesicles. (A) Representative particle size distribution of EVs isolated by ultracentrifugation (UC), co-precipitation (Q), and immunoaffinity capture (M), as determined by nanoparticle tracking analysis (NTA) (n = 4). (B) Representative cryo-transmission electron microscopy (cryo-TEM) images of EVs obtained using each isolation method (UC, Q, M). Scale bars are indicated on each image. (C) Mean protein yield (µg) obtained from 1 mL of saliva per isolation method. p < 0.01 is denoted by **. (D) EV purity index calculated as the ratio of particle number to total protein content. (E) Western blot analysis in WS samples and EV fractions isolated by UC, Q, and M.

**Table 1.** Salivary EV metrics by isolation method.

| EV isolation method | EV concentration (particle/ml) | EV mean size (nm) | EV mode (nm) | Protein concentration (µg/ml) | Purity |
| --- | --- | --- | --- | --- | --- |
| UC | 4,45E+08 ± 2,74E+08 | 253,3 ± 18,5 | 180,4 ± 42,8 | 76,0 ± 76,6 | 1,87E+07 ± 2,45E+06 |
| Q | 1,93E+08 ± 4,94E+07 | 224,1 ± 15,7 | 146,6 ± 24,1 | 106,6 ± 71,0 | 3,24E+06 ± 1,70E+05 |
| M | 1,48E+09 ± 5,24E+08 | 83,6 ± 10,8 | 62,4 ± 6,1 | 152,8 ± 26,5 | 1,04E+07 ± 1,96E+06 |

Cryo-EM confirmed the presence of vesicular structures in all preparations. UC samples occasionally showed vesicle-in-vesicle structures, Q samples exhibited some agglutination with PEG-like residues, and M isolates were well dispersed, though visualization was partly obscured by antibody-coated beads (Fig. 2B).

WB analysis confirmed enrichment of EV markers (CD63, CD9, TSG101) in the EVs compared to WS. Albumin was abundant in WS, faint in UC samples, more pronounced in Q, and negligible in M, consistent with their respective purity indices (Fig. 2E, uncropped membranes on Fig. S1).

Overall, all three methods successfully recovered EV-like particles, with distinct profiles in yield, size distribution, and co-isolated proteins (Table 1, Fig. 2).

### Proteomic analysis

Label-free quantitative proteomics (LC-MS/MS) was performed on WS and EVs isolated by UC, Q, and M (n = 3 per method). Using stringent identification criteria (≥2 unique peptides, detection in ≥2 replicates), we identified an average of 781 proteins in WS, 974 in UC-derived EVs, 902 in Q-derived EVs, and 449 in M-derived EVs (Fig. 3A).

**Figure 3.**
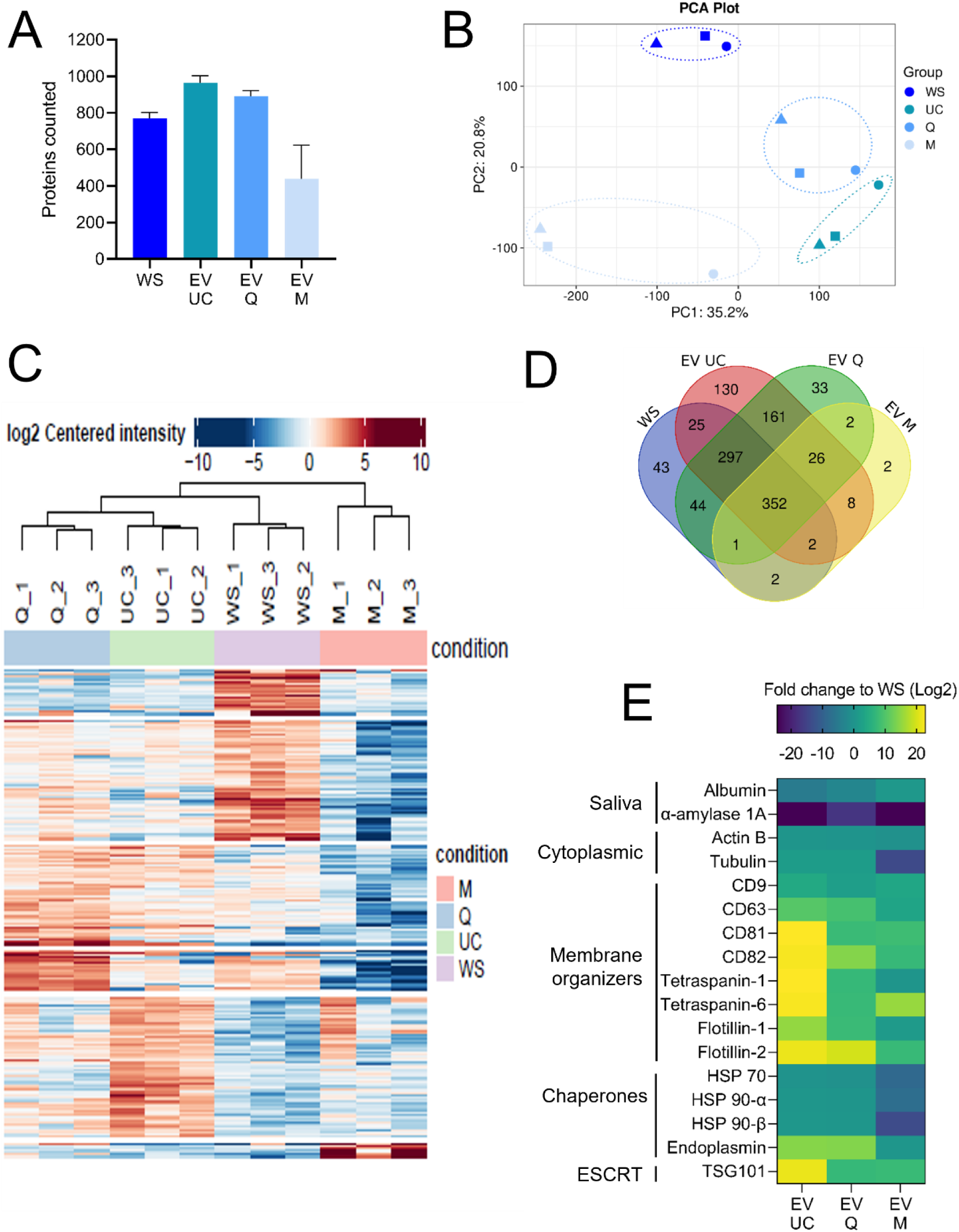
Proteomic analysis of whole saliva and EV fractions. (A) Mean number of proteins identified in whole saliva (WS), and in EVs isolated by ultracentrifugation (UC), co-precipitation (Q), and immunoaffinity (M). Proteins included had at least two peptides and were detected in a minimum of two out of three replicates. (B) Principal component analysis (PCA) of proteomic profiles showing separation between WS and EV samples based on variance in protein expression. (C) Hierarchical clustering heatmap of all samples based on normalized protein abundance. (D) Venn diagram illustrating the overlap of identified proteins between WS (blue), EV UC (red), EV Q (green), and EV M (yellow). (E) Heatmap representing fold change (relative to WS) in EV UC, Q, and M for a subset of proteins commonly reported as salivary or EV-associated markers.

Principal component analysis (PCA) indicated that isolation method was the main source of proteomic variation (Fig. 3B). UC and Q isolates clustered closely, while M-derived EVs and WS formed separate groups. Hierarchical clustering of 230 significantly differentially expressed proteins confirmed this pattern (Fig. 3C and Fig. S2). Across conditions, 352 proteins were shared between WS and EVs, while 362 were unique to EV preparations. UC and Q shared 161 proteins, compared with only 36 shared between M and the other methods. Both UC and Q also showed broader overlap with WS (297 shared proteins) (Fig. 3D).

Comparative abundance analysis revealed consistent enrichment of EV-associated proteins in all isolates compared to WS (Fig. 3E). UC and Q showed the strongest enrichment, whereas M isolates had lower overall protein abundance, consistent with reduced yield. Salivary contaminants such as albumin and α-amylase were depleted across EV samples relative to WS.

Functional enrichment (KEGG) revealed method-specific signatures. UC and Q isolates were enriched in proteasome-related and intracellular pathways (Fig. 4A, 4C), while M isolates showed enrichment in lysosomal and vesicle trafficking proteins, as well as salivary secretion components (Fig. 4E). Gene Ontology (cellular component) analysis confirmed enrichment of typical EV terms across all methods, though statistical strength was higher in UC and Q (adjusted p-values 1e–26 to 1e–17) than in M (∼1e–9) (Fig. 4B, 4D, 4F). These results indicate that UC and Q yield broader and more complex proteomic profiles, while M isolates produce a more restricted but focused protein set.

**Figure 4.**
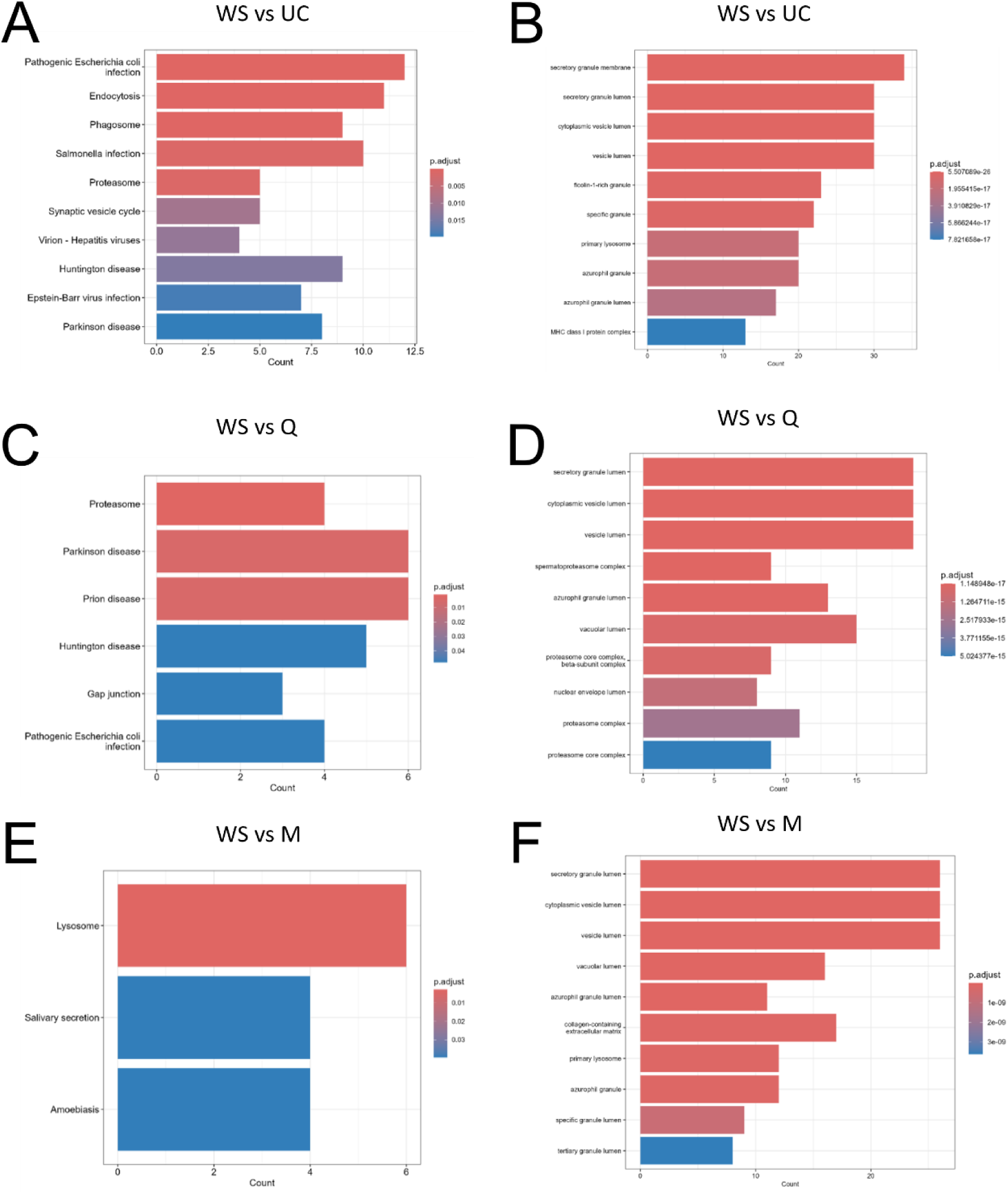
Functional enrichment analysis of differentially abundant proteins between whole saliva and EV fractions. Bar plots illustrate the results of functional enrichment analyses performed using the ClusterProfiler R package (v4.10.1) on proteins differentially abundant between WS and each EV isolation method. (A, C, E) KEGG pathway enrichment for WS vs. UC, WS vs. Q, and WS vs. M, respectively. (B, D, F) Gene Ontology (GO) enrichment (cellular component category) for WS vs. UC, WS vs. Q, and WS vs. M, respectively. The x-axis indicates the number of proteins associated with each enriched pathway or cellular component. The y-axis lists the top enriched terms. Bar color reflects the adjusted p-value (Benjamini-Hochberg), with a redder color indicating stronger statistical significance.

### Global transcriptomic profiling of miRNAs

To evaluate the impact of isolation method on RNA cargo, we performed small RNA sequencing on WS and EVs obtained by UC, Q, and M. Total small RNA was extracted with the Qiagen miRNeasy kit and quantified using a Qubit microRNA assay. M-derived samples yielded lower miRNA amounts, consistent with their more selective enrichment (Fig. 5A). Sequencing libraries were prepared with the Illumina TruSeq small RNA kit and analysed on a NextSeq 500. After quality filtering and adapter trimming, only miRNAs detected in all three biological replicates in at least one condition were retained.

**Figure 5.**
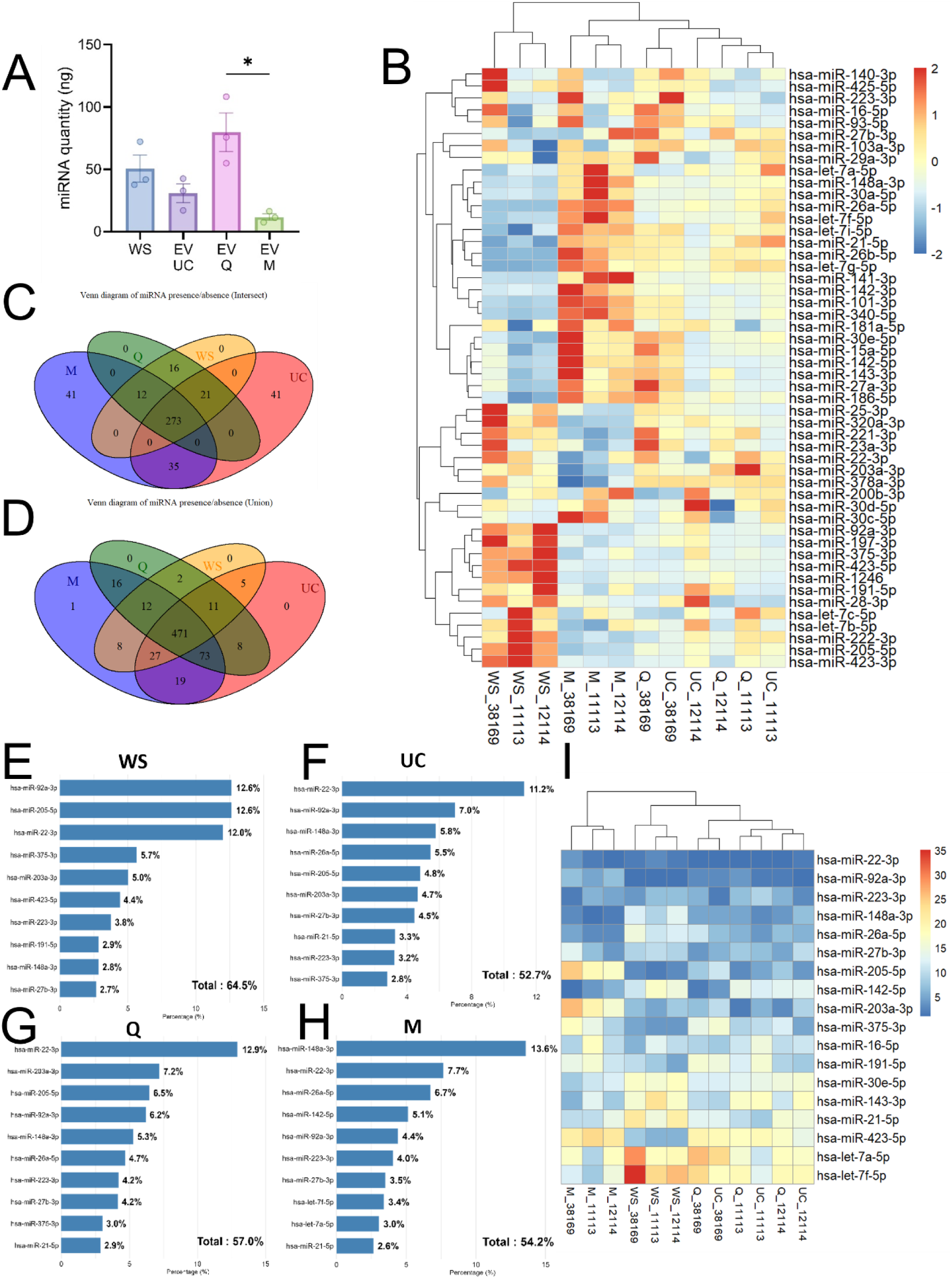
MicroRNA profiling of whole saliva and EV isolates. (A) Total miRNA quantity measured in whole saliva (WS) and EV samples isolated by ultracentrifugation (UC), precipitation (Q), and immunoaffinity capture (M). (B) Heatmap of the top 50 most abundant miRNAs based on counts per million (CPM). Values are Z-score normalized (−2 to +2) to highlight relative abundance across conditions (WS, UC, Q, M). Color gradient represents abundance from low (blue) to high (red). (C–D) Venn diagrams showing the overlap of miRNA species detected in WS, UC, Q, and M. (C) Union: miRNAs detected in at least one replicate per condition. (D) Intersection: miRNAs detected in all three replicates per condition. Colors denote conditions (WS: yellow, UC: red, Q: green, M: purple). (E–H) Bar plots of the top 10 most abundant miRNAs in each representative sample (WS_11113, UC_11113, Q_11113, M_11113). Relative abundance is expressed as percentage of total CPM. (I) Rank-based heatmap of miRNAs appearing in the top 10 positions (by CPM) in at least one sample. Color gradient reflects rank. X-axis represents individual replicates across conditions; Y-axis lists the top-ranked miRNAs.

Hierarchical clustering of the 50 most abundant miRNAs showed clear differences between WS and EVs (Fig. 5B). WS samples formed a distinct group, while EV isolates shared closer expression patterns. UC and Q clustered by donor, indicating high similarity between the two methods, whereas M-derived EVs clustered by isolation method, suggesting a more distinct miRNA cargo.

Presence–absence analysis indicated that no miRNAs were uniquely detected in WS, while multiple miRNAs were specific to EV fractions (Fig. 5C–D). The top 10 most abundant miRNAs accounted for ∼53–65% of total reads, underscoring the dominance of a few highly expressed species (Fig. 5E–H, Fig. S3). Among the consistently enriched miRNAs across EVs were hsa-miR-22-3p, hsa-miR-92a-3p, and hsa-miR-223-3p (Fig. 5I).

Overall, EV-associated miRNA profiles differed markedly from WS and varied across isolation methods, indicating that method choice influences both yield and composition.

### Differential expression analysis of miRNAs

To assess how isolation method influences EV-associated miRNA cargo, we compared expression profiles of EVs obtained by UC, Q, and M against WS. Only miRNAs detected in all three biological replicates of a condition were retained, yielding 731 miRNAs across comparisons.

Volcano plots highlighted significant differences in abundance between WS and EV fractions (Fig. 6A). The M method displayed the broadest range of significantly downregulated miRNAs (log₂FC < 0), consistent with selective enrichment of EV-associated species.

**Figure 6.**
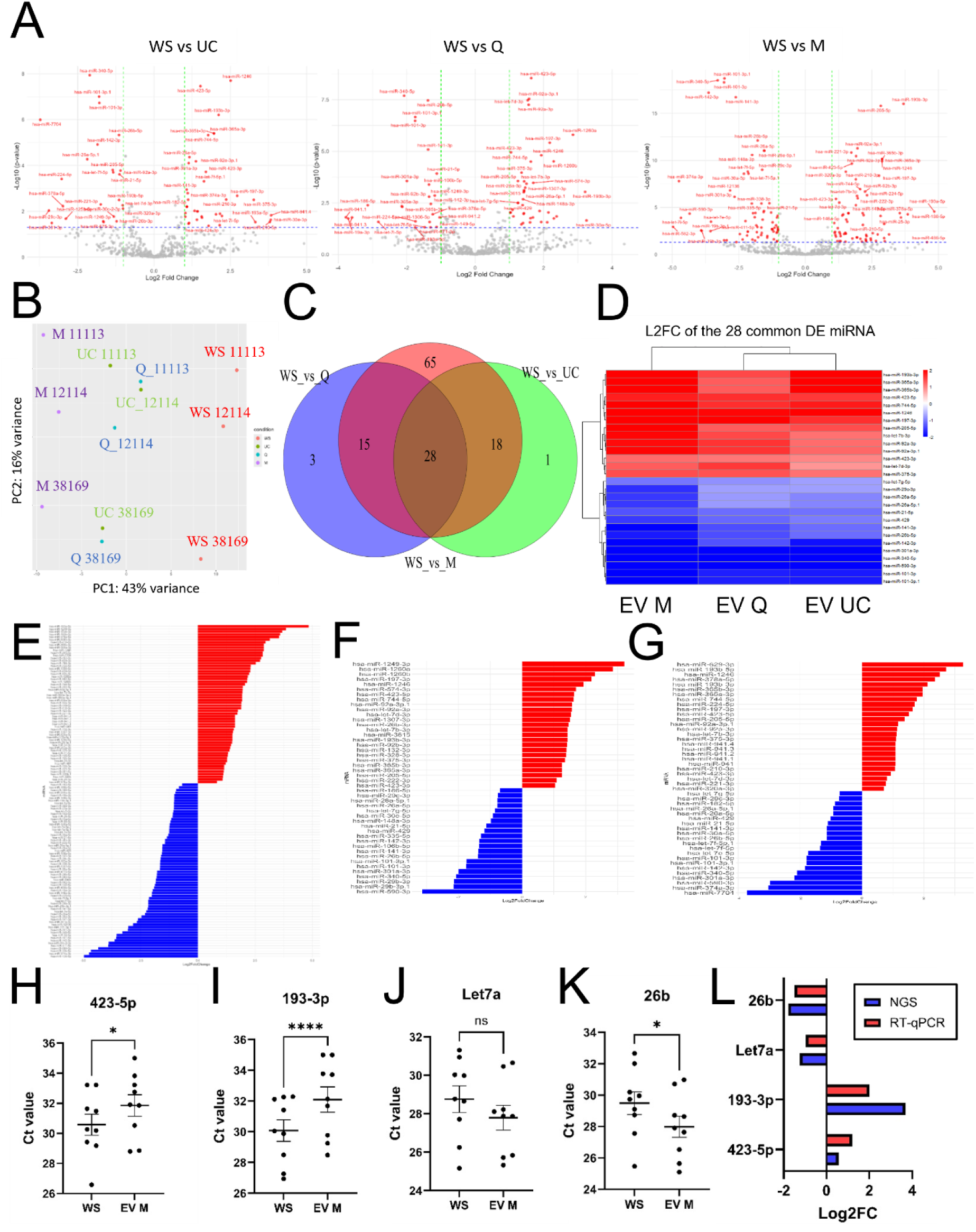
Differential expression analysis of salivary miRNAs across EV isolation methods. (A) Volcano plots showing differentially expressed miRNAs between whole saliva (WS) and EV isolates: UC, Q, and M. The x-axis represents log₂ fold change (log₂FC), and the y-axis indicates statistical significance as−log₁₀(p-value). Red dots mark significantly regulated miRNAs (adjusted p-value < 0.05; |log₂FC| > 1). Right shift (positive log₂FC) indicates upregulation in WS; left shift (negative log₂FC) indicates enrichment in EVs. Grey dots represent non-significant changes. Plots were generated with DESeq2 (v1.42.1) and ggplot2 (v3.5.1). (B) Principal component analysis (PCA) of global miRNA profiles from WS and EV isolates (UC, Q, M), each with three biological replicates. PC1 and PC2 account for 43% and 16% of total variance, respectively. Analysis was performed using DESeq2. (C) Venn diagram of differentially expressed miRNAs (adjusted p < 0.05) in WS vs. UC (green), WS vs. Q (purple), and WS vs. M (red). Overlaps represent miRNAs commonly regulated across methods. Diagram generated using the R package *VennDiagram* (v1.7.3). (D) Heatmap of log₂ fold changes for the 28 miRNAs significantly and commonly regulated across all three comparisons (WS vs. UC, Q, M). Rows represent miRNAs; columns represent each comparison. Colors range from red (upregulated in WS) to blue (upregulated in EVs). Heatmap created using *pheatmap* (v1.0.12); log₂FC values scaled from −2 to +2. (E–G) Bar plots showing log₂ fold changes of all significantly regulated miRNAs in: (E) WS vs. M (n = 47 miRNAs), (F) WS vs. Q (n = 46 miRNAs), (G) WS vs. UC (n = 128 miRNAs). Red bars indicate miRNAs enriched in WS; blue bars indicate those enriched in EVs. Bar plots created using *ggplot2* (v3.5.1). (H-K) Ct values of miRNAs in WS compared to EV M measured by RT-qPCR. miRNA hsa-423-5p (H), hsa-193-3p (I), hsa-let7a (J) and hsa-26b (K) are represented with mean and SEM values (n=9). *: p < 0.05; ****: p < 0.0001. (L) Comparison of Log2FC calculated from NGS and from RT-qPCR datasets.

Principal component analysis further emphasized method effects (Fig. 6B). M-derived EVs separated clearly from WS along PC1, which explained 43% of total variance, while UC and Q clustered closely, suggesting similar EV-associated miRNA populations. PC2 (16% of variance) reflected donor-related variation across all groups. These patterns indicate that immunoaffinity capture yields a distinct miRNA profile, whereas UC and Q capture more heterogeneous populations.

DESeq2 analysis (FDR < 0.05) identified 47, 46, and 128 differentially expressed miRNAs in UC, Q, and M compared to WS, respectively (Fig. S4). A Venn diagram revealed 28 miRNAs consistently altered across all EV methods, while 65 were unique to M, versus 3 for Q and 1 for UC (Fig. 6C).

Heatmaps confirmed these trends, with several miRNAs consistently enriched in EVs (e.g., hsa-let-7f-5p, hsa-let-7a-5p, hsa-miR-21-5p, hsa-miR-143-3p), while others were more abundant in WS (e.g., hsa-miR-423-5p, hsa-miR-375-3p) (Fig. 6E–G).

Overall, these results demonstrate that EV isolation method substantially shapes the detectable miRNA landscape, influencing both overlap with WS and the recovery of potentially informative EV-specific miRNAs.

### RT-qPCR validation

To validate NGS-derived differential expression patterns, RT-qPCR was performed on EVs isolated by the M method, which showed the most distinct miRNA profile and the largest number of unique differentially expressed species. Four candidates were selected: two enriched in WS (hsa-miR-423-5p, hsa-miR-193a-3p) and two enriched in EVs (hsa-miR-26b-5p, hsa-let-7a-5p), based on NGS fold-change values.

RT-qPCR was carried out on nine biological replicates. Despite inter-individual variation, expression patterns were consistent with sequencing results: hsa-miR-423-5p and hsa-miR-193a-3p were higher in WS (Fig. 6H–I), while hsa-miR-26b-5p and hsa-let-7a-5p were enriched in EVs (Fig. 6J–K). Calculated log₂ fold-change values (−ΔCt, WS as reference) confirmed both direction and magnitude of enrichment observed by NGS (Fig. 6L).

These results support the robustness of the sequencing dataset and confirm that EV isolation influences detectable miRNA abundance.

### Functional enrichment analysis of validated and predicted target genes of differentially expressed miRNAs

To explore potential biological relevance, functional enrichment was performed on validated and predicted targets of differentially expressed miRNAs. Analyses focused on the 28 miRNAs commonly altered across all EV methods and the 65 miRNAs uniquely identified in M-derived EVs.

For the shared set, enrichment of >15,000 validated and >14,000 predicted targets was observed, while M-specific miRNAs were associated with slightly larger target pools (>16,000 validated, >17,000 predicted).

Gene Ontology (cellular component) analysis indicated consistent associations with neuronal structures, including postsynaptic density, synapses, and axons (Fig. 7A; Fig. S5–S6). KEGG pathway analysis highlighted enrichment in axon guidance, autophagy, MAPK signalling, and several neurodegeneration-related pathways for the shared miRNAs (Fig. 7B). M-specific miRNAs showed overlapping but broader patterns, with additional enrichment in pathways such as cell cycle regulation, focal adhesion, and proteoglycans in cancer (Fig. S7).

**Figure 7.**
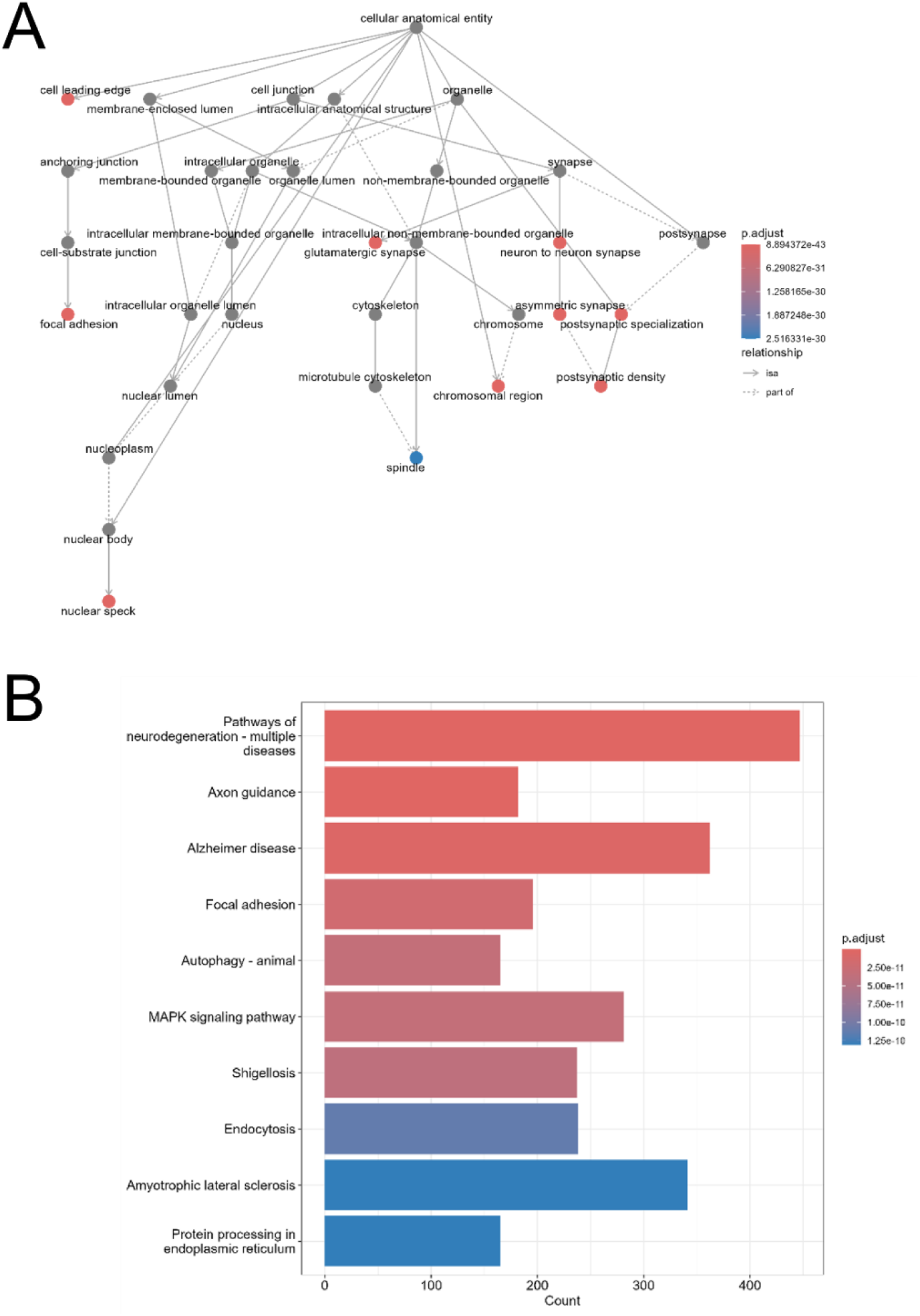
Functional enrichment analysis of target genes regulated by common salivary miRNAs. (A) Gene Ontology (GO) enrichment network for cellular component (CC) terms associated with 15,555 predicted target genes of the 28 commonly regulated miRNAs across all EV isolation methods (WS vs. M, Q, UC). Nodes represent GO terms; solid arrows indicate hierarchical subcategories; dashed arrows show associative links. Node color reflects adjusted p-value: red for most significant, blue for least, and grey for non-significant terms. Analysis and visualization performed using the *clusterProfiler* R package (v4.10.1). (B) KEGG pathway enrichment of the same 15,555 target genes. The x-axis shows the number of genes per pathway; the y-axis lists enriched KEGG pathways. Bar color gradient reflects statistical significance (adjusted p-value), with red indicating higher significance and blue indicating lower. Analysis was conducted with *clusterProfiler* (v4.10.1) using the KEGG database (https://www.genome.jp/kegg/).

These results indicate that both shared and method-specific salivary EV miRNAs are predicted to regulate genes involved in neuronal and signalling processes. However, the enrichment patterns differ depending on the isolation method, reflecting differences in captured EV populations.

## Discussion

In this study, we performed a systematic comparison of three commonly used EV isolation methods: ultracentrifugation (UC), PEG-based precipitation (Q), and immunoaffinity capture (M); applied to human saliva. By integrating proteomic and transcriptomic analyses, we show that isolation strategy not only affects vesicle yield and purity, but also strongly influences the qualitative and quantitative composition of salivary EV cargo. These findings emphasize that methodological variability can shape biological readouts and must therefore be carefully considered in biomarker discovery and translational EV research. To support method selection, we summarize the advantages and downstream compatibility of each protocol in Table 2.

**Table 2.** Practical aspects and downstream analysis compatibility of each isolation method.

|  | Ultracentrifugation | Coprecipitation | Immuno-capture |
| --- | --- | --- | --- |
| Mechanism of separation | Size and density | Surface charge. Use of a cationic polymer to coprecipitate EVs | Specific binding of antibodies against EV markers tetraspanins CD9, CD63 and CD81 coated on magnetic beads |
|  | Centrifugation at 100,000 xG |  |  |
| Specificity | ** | * | *** |
| Convenience/practicability | * | *** | ** |
| Processing time | *** | * | * |
| Hands-on time | *** | * | ** |
| Cost | ** | * | *** |
| Scalability | * | *** | * |
| Downstream Analysis Compatibility |  |  |  |
| NTA | *** | *** | * |
| Cryo-EM | *** | ** | * |
| Protein concentration | *** | *** | ** |
| Western Blot | *** | *** | *** |
| RT-qPCR | *** | *** | *** |
| Proteomics | *** | *** | * |
| NGS | *** | *** | *** |

### EV isolation method shapes the proteomic landscape

Our proteomic analysis showed that UC and Q recovered broader and more diverse protein repertoires than M. Despite differences in yield and particle size, UC and Q clustered closely in PCA, indicating that both methods capture a heterogeneous mix of EV subtypes along with co-isolated non-vesicular proteins. These isolates were enriched in cytosolic proteins, proteasome complexes, and pathways linked to neurodegeneration and intracellular trafficking, consistent with the molecular complexity of saliva. Similar patterns have been observed in salivary EV studies where broader enrichment strategies improved detection of low-abundance proteins relevant for biomarker discovery ^26^, as well as in plasma EV proteomics where UC and precipitation-based methods achieved high proteome coverage but introduced contamination from abundant proteins such as albumin and apolipoproteins ^49,50^.

By contrast, immunoaffinity capture (M), which targets tetraspanins (CD9, CD63, CD81), yielded a more restrictive proteome enriched in lysosomal and vesicle trafficking proteins. This pattern is consistent with the isolation of a narrower small EV subpopulation, likely of endosomal origin and enriched in epithelial or salivary gland–derived vesicles, as previously described for tetraspanin-positive salivary EVs ^51^. Although all three methods were enriched in canonical EV-associated terms in GO analysis, UC and Q showed greater statistical significance, reflecting higher proteomic complexity rather than differences in EV quality ^52^.

These results enlighten recent work by Kawano et al.^53^, who described distinct molecular profiles in salivary EV subtypes, and by Sandira et al. ^54^, who showed that technical variability introduced by isolation methods can confound biological interpretation. Collectively, our results highlight a practical dichotomy: UC and Q are better suited for exploratory proteomics and broad-spectrum biomarker discovery, whereas immunoaffinity isolation provides higher purity and reproducibility for targeted applications where background interference must be minimized.

### Method-dependent variation in EV miRNA landscapes

Small RNA sequencing revealed that EV-associated miRNA profiles differed substantially both from WS and across isolation methods. As expected, several highly abundant salivary miRNAs were depleted in EV fractions, consistent with previous reports showing that vesicle-encapsulated RNAs represent a selective subset rather than a simple reflection of cell-free biofluid content ^55,56^.

PCA indicated that isolation method was the dominant source of variation, exceeding donor-dependent effects. UC and Q samples clustered closely, suggesting they capture similar vesicle populations, while M-derived EVs formed a distinct cluster. Importantly, this separation reflects selective enrichment of a specific EV subpopulation by immunoaffinity capture, rather than improved inter-individual reproducibility. This subtlety is in line with observations in serum by Buschmann et al.^57^ who demonstrated that isolation strategy strongly influences small RNA landscapes, particularly with respect to low-abundance species.

Differential expression analysis further highlighted method-specific signatures. Twenty-eight miRNAs were consistently altered across all three EV methods compared to WS, suggesting a core EV-associated signature. In contrast, M-derived EVs contained sixty-five unique differentially expressed miRNAs, compared to only a few in UC or Q. This pattern suggests that immunoaffinity capture facilitates detection of low-abundance or selectively packaged miRNAs that may be missed by broader enrichment strategies. These observations reinforce our previous observations in saliva ^43^ as well as other study by Llorens-Revull et al. in plasma ^35^, where isolation method determined both the diversity and detectability of miRNAs.

Importantly, NGS-derived differences were confirmed by RT-qPCR for selected candidates (e.g., hsa-let-7a-5p, hsa-miR-26b-5p, hsa-miR-423-5p, hsa-miR-193a-3p), supporting the robustness of our dataset and aligning with best practices in EV biomarker studies, which emphasize orthogonal validation for translational reliability.

Together, these results indicate that while UC and Q provide broader coverage of salivary EV miRNAs, immunoaffinity capture shows a more distinct transcriptomic profile. This trade-off has direct implications for biomarker discovery: broad methods may maximize sensitivity for exploratory studies, whereas selective capture may improve detection of biologically specialized miRNAs with potential mechanistic or diagnostic value.

### Functional implications: neurodegeneration-linked miRNAs in salivary EVs

Functional enrichment of validated and predicted targets from differentially expressed miRNAs revealed strong associations with neuronal structure and signalling. GO terms such as postsynaptic density, axon, and synapse were consistently overrepresented, highlighting processes central to synaptic plasticity and vulnerable in neurodegenerative disease. KEGG pathway analysis further identified enrichment in Alzheimer’s disease, amyotrophic lateral sclerosis (ALS), and MAPK signalling, in line with previous studies reporting that plasma-and CSF-derived EVs carry neuronally relevant cargo reflective of CNS pathophysiology ^58,59^. Our findings extend this concept to saliva, an easily accessible biofluid suitable for repeated sampling and clinical translational studies.

The 28 miRNAs consistently altered across all EV isolation methods were enriched for neuronal and synaptic pathways, suggesting a conserved salivary EV miRNA signature with potential biomarker relevance. These findings expand on prior work by Rastogi et al. ^19^, who reported that salivary EVs reflect systemic molecular alterations in neurodegenerative contexts, supporting their biomarker potential. Notably, the 65 miRNAs uniquely detected in M-derived EVs mapped to additional pathways including cell cycle regulation, Hippo signalling, and proteoglycans in cancer; networks implicated in neural stem cell homeostasis, apoptosis, and neuroinflammation. This selective enrichment may reflect the ability of tetraspanin-targeted capture to isolate specialized EV subpopulations, as observed in neural-derived EV studies ^60,61^.

Beyond neurological pathways, recent studies emphasize the broader biological responsiveness of salivary EV miRNAs. Kazanopoulos et al. ^62^ identified over 1,400 distinct salivary EV miRNAs and demonstrated time-dependent changes during orthodontic treatment, underscoring that salivary EVs can capture dynamic remodeling processes. Together with our results, these observations reinforce the potential of salivary EVs as information-rich sources for monitoring both systemic and disease-specific biological states.

Our observations are also consistent with recent translational work. Ryu et al. ^17^ showed that salivary EV-derived miR-485-3p correlated with cerebral amyloid-β burden and predicted PET positivity in Alzheimer’s disease patients, while Malaguarnera et al. ^63^ further highlighted miR-485-3p among leading salivary EV miRNAs with diagnostic relevance in neurodegenerative diseases, supporting its inclusion in candidate biomarker panels. Collectively, these studies highlight that salivary EV miRNAs can provide non-invasive readouts of neuronal processes and complement established CSF or plasma biomarkers.

### Strategic recommendations for salivary EV isolation

Our results reinforce that no single isolation method is equivalent; instead, selection should be guided by research objectives and analytical priorities. For exploratory proteomics and discovery-driven studies, UC and Q provide broader coverage of the salivary EV proteome and miRNA repertoire. Their ability to recover heterogeneous vesicle populations increases sensitivity for detecting diverse biomolecules, although this comes at the cost of co-isolating non-vesicular contaminants. For targeted transcriptomics, biomarker validation, or clinical assay development, immunoaffinity capture (M) offers greater selectivity for CD9/CD63/CD81-positive EVs, reducing background proteins and yielding distinct miRNA signatures. While not necessarily more reproducible across individuals, its specificity makes it suitable when analytical sensitivity and purity are critical.

In translational pipelines, combining orthogonal approaches or cross-validating results across multiple methods can mitigate isolation-specific biases and strengthen reproducibility. This layered strategy may be particularly valuable when moving from biomarker discovery to clinical validation.

These recommendations align with the MISEV2023 guidelines ^44^, which emphasize that isolation strategy must be tailored to the biofluid, analyte of interest, and downstream platform. Failure to account for these factors can introduce technical variability that exceeds biological differences, as highlighted by Sandira et al. ^54^. Moving forward, harmonization of protocols and transparent reporting will be essential to enable cross-study comparability and accelerate clinical translation of salivary EV biomarkers.

## Conclusion

This study provides the first systematic multi-omics comparison of ultracentrifugation, PEG-based precipitation, and immunoaffinity capture for isolating salivary EVs. We show that isolation strategy profoundly affects vesicle yield, purity, and molecular cargo, including both proteins and miRNAs. UC and Q recover broader and more heterogeneous proteomic and transcriptomic profiles, supporting their use in discovery-driven applications. In contrast, immunoaffinity capture enriches for a more selective EV subset with distinct miRNA signatures, some linked to neuronal and neurodegenerative pathways. These findings highlight that isolation choice must be aligned with experimental objectives and that method-specific biases should be carefully considered in biomarker development.

### Limitations and future directions

Several limitations should be acknowledged. First, the omics analyses were conducted on a limited number of biological replicates (n=3), restricting statistical power. Larger, more diverse cohorts are needed to validate isolation-specific signatures and capture inter-individual variability. Second, RT-qPCR validation was performed only on M-derived EVs; extending validation to UC-and Q-derived EVs will help confirm the robustness of observed transcriptomic differences. Third, functional enrichment was based on computational predictions and validated target databases, which require experimental follow-up to confirm biological relevance, particularly for pathways linked to neurodegeneration.

Methodologically, size exclusion chromatography (SEC) was not included despite its widespread use for EV purification ^64,65^. However, SEC was not included in this study due to practical and methodological constraints. SEC offers high purity but requires larger input volumes and is sensitive to saliva’s viscosity and mucin content, which can reduce yield and column performance ^66,67^. Our study instead focused on methods commonly applied in translational workflows (UC, Q, M). Future studies should include SEC and emerging isolation technologies to build a more comprehensive framework for salivary EV research.

Beyond these analytical and methodological considerations, future work should also integrate advanced single-particle characterization approaches to achieve a more robust and quantitative assessment of EV properties. Conventional bulk techniques such as NTA and Western blotting provide limited information on vesicle heterogeneity and marker distribution. In contrast, nano-flow cytometry now enables simultaneous measurement of EV size, concentration, and surface marker expression at the single-particle level, representing an emerging gold standard for EV characterization ^68^. Applying it to saliva-derived EVs would allow determination of the proportion of tetraspanin-positive vesicles (CD9⁺, CD63⁺, CD81⁺) and quantification of the fraction of lipid bilayer-containing particles, using fluorescent probes such as CMDR or ExoBrite dyes. Implementing such analyses across different isolation methods could provide critical quantitative insights into EV purity, composition, and biological relevance, supporting the establishment of standardized benchmarks for salivary EV characterization.

Moving forward, harmonizing isolation protocols, implementing robust quality control metrics, and improving automation will be critical to reduce technical variability and enable reproducibility across studies. Such standardization will accelerate the integration of salivary EVs into biomarker pipelines, supporting their potential as a scalable, non-invasive tool for precision diagnostics, including monitoring of neurodegenerative disease.

## Aknowledgements

The authors would like to thank Dr. Marie Morille, Institut Charles Gerhardt Montpellier, UMR 5253 CNRS-ENSCM-UM, Equipe Matériaux Avancés pour la Catalyse et la Santé, Montpellier, for her valuable technical assistance for nanoparticles tracking analysis and Dr. Josephine Lai Kee Him and Mrs. Aurelie Ancelin, Centre de Biochimie Structurale, Université de Montpellier, CNRS, INSERM, for their Cryo-EM analysis. Mass spectrometry was carried out using the facilities of the Montpellier Proteomics Platform (PPM, BioCampus Montpellier). We also thank all the study participants.

## Author contributions

Study concept and design: F.M. and M.K. Acquisition, analysis, interpretation of data: All authors. Drafting of the manuscript and critical revision of the manuscript for important intellectual content: F.M. and M.K. Statistical analysis: E.S. and M.K. Funding acquisition: F.M. All authors critically reviewed and edited the manuscript.

## Data availability statement

The authors declare that all relevant data have been provided within the manuscript and its supporting information files.

## Funding

This project was supported by Centre National de la Recherche Scientifique (CNRS) and ALCEN.

## Competing interests statement

The authors declare that there are no conflicts of interest.

## Notes

### Competing Interest Statement

The authors have declared no competing interest.

