## Supplementary Material for "Comparison of extracellular vesicles isolation methods reveals method-dependent protein and miRNA profiles in saliva"

[1]Sys2Diag UMR9005 CNRS/ALCEN, Cap Gamma, Parc Euromédecine, 1682 rue de la Valsière, CS 40182, 34184, Montpellier, CEDEX 4, France.

***Corresponding author:**

### Figure S1.

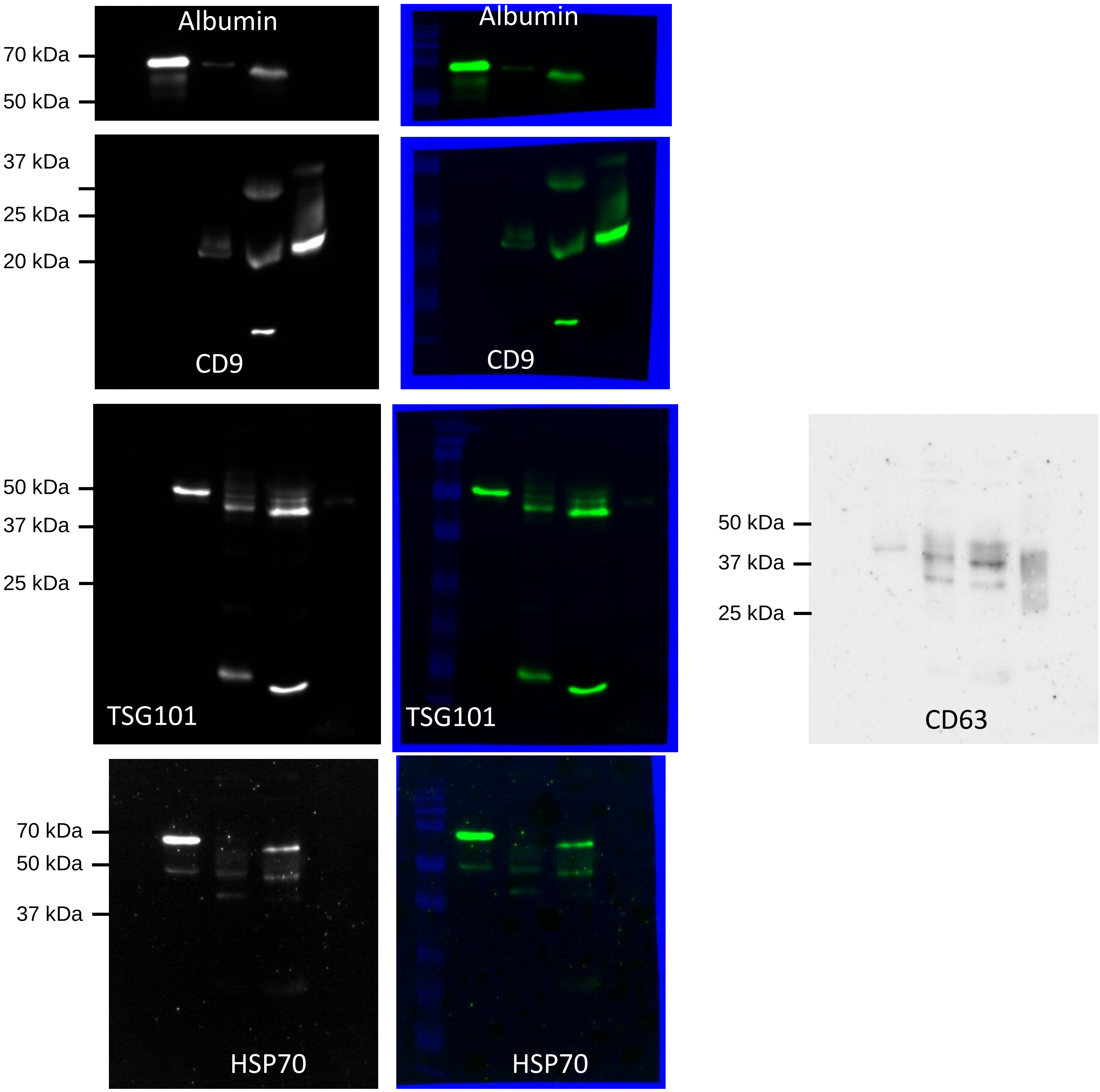

**Figure S1: Full-length and cut membranes of all immunoblots related to Fig. 2E.** Uncropped Western blots corresponding to Fig.2 (E). Molecular Weight are indicated on the left side of the membrane. Some membranes (Albumin and CD9) were cut prior to hybridization with antibodies.

Figure S2.

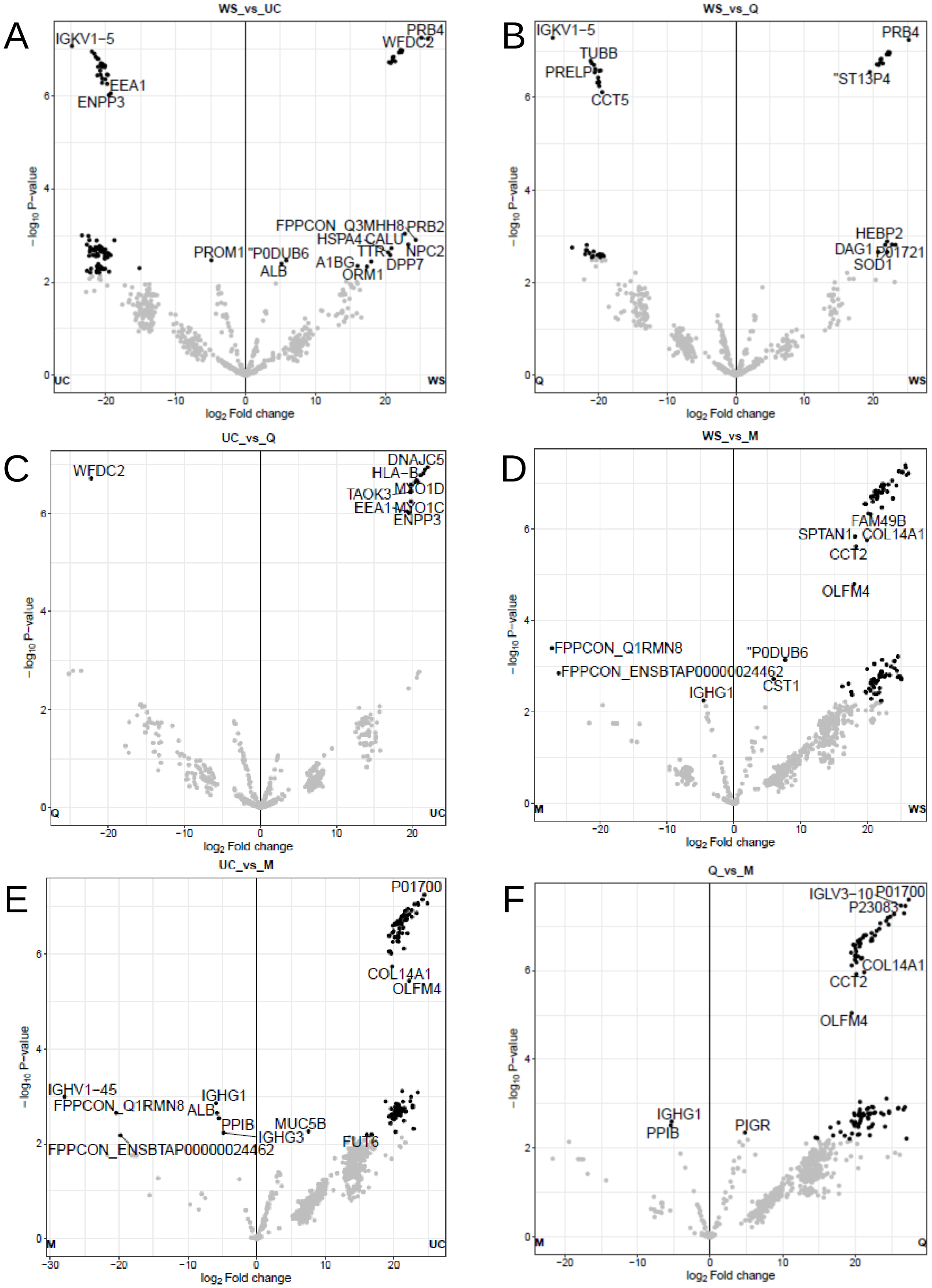

**Figure S2: Proteomic profiles of salivary EVs and whole saliva.** Volcano plots depicting differentially expressed proteins between whole saliva (WS) and salivary extracellular vesicle (EV) isolated by ultracentrifugation (UC), co-precipitation (Q), or immunoaffinity capture (M). Comparisons include: (A) WS vs. UC, (B) WS vs. Q, (C) WS vs. M, (D) UC vs. Q, (E) UC vs. M, and (F) Q vs. M. The x-axis indicates $\log_2$ fold change ($\log_2FC$); the y-axis represents $-\log_{10}$(p-value). Proteins meeting significance criteria (adjusted p-value $< 0.05$ and $|\log_2FC| > 1$) are shown in black. Non-significant proteins are shown in grey. Plots were generated using *LFQ-Analyst*.

Figure S3.

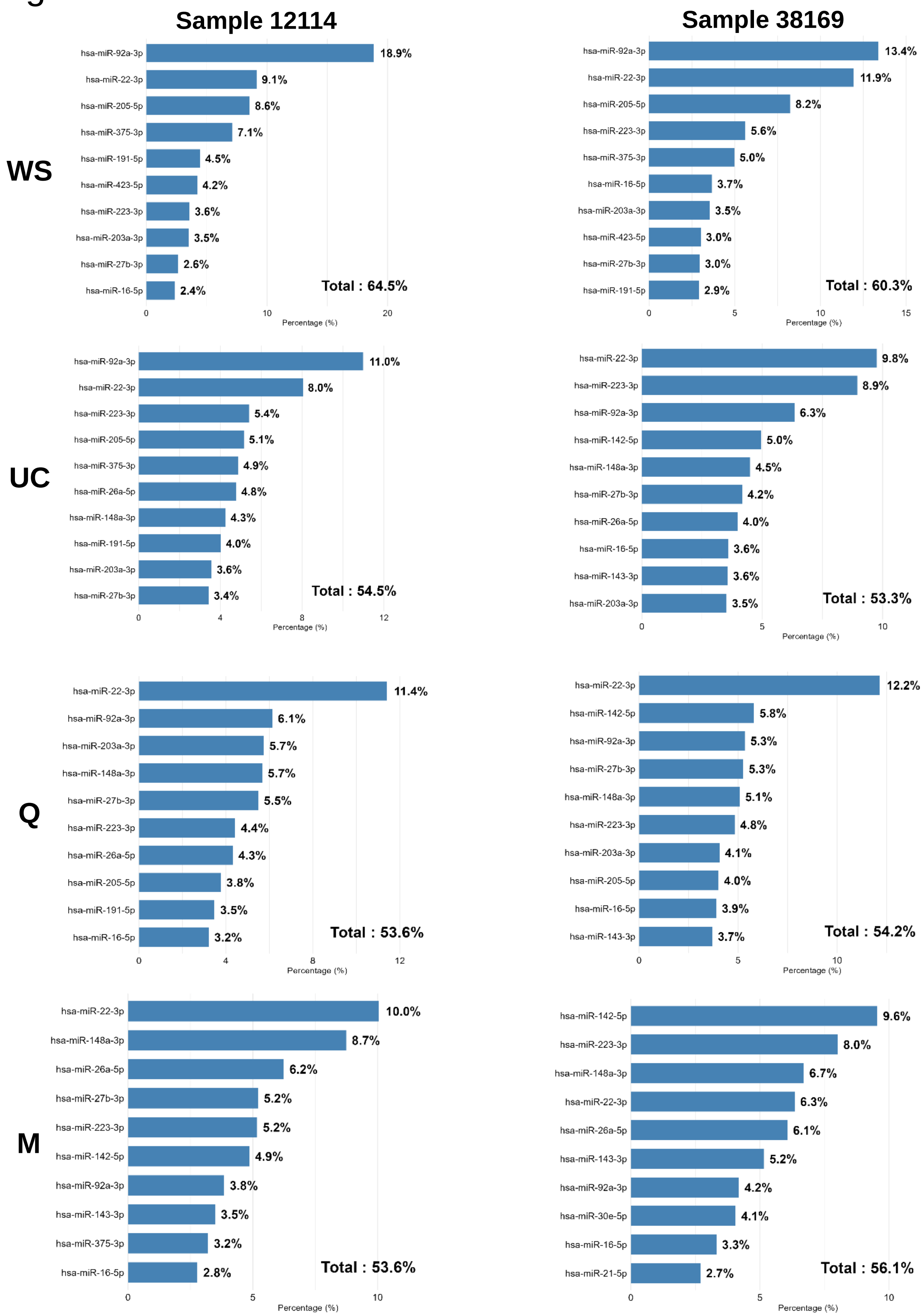

**Figure S3. Top 10 most abundant miRNAs in individual samples.** Bar plots displaying the top 10 most abundant microRNAs (miRNAs) in two individual donors: sample 12114 (left) and sample 38169 (right), across conditions: whole saliva (WS), ultracentrifugation (UC), co-precipitation (Q), and immunoaffinity (M), respectively. Relative abundance is shown as the percentage of total counts per million (CPM). Graphs were generated using the R package ggplot2 (version 3.5.1).

### Figure S4.

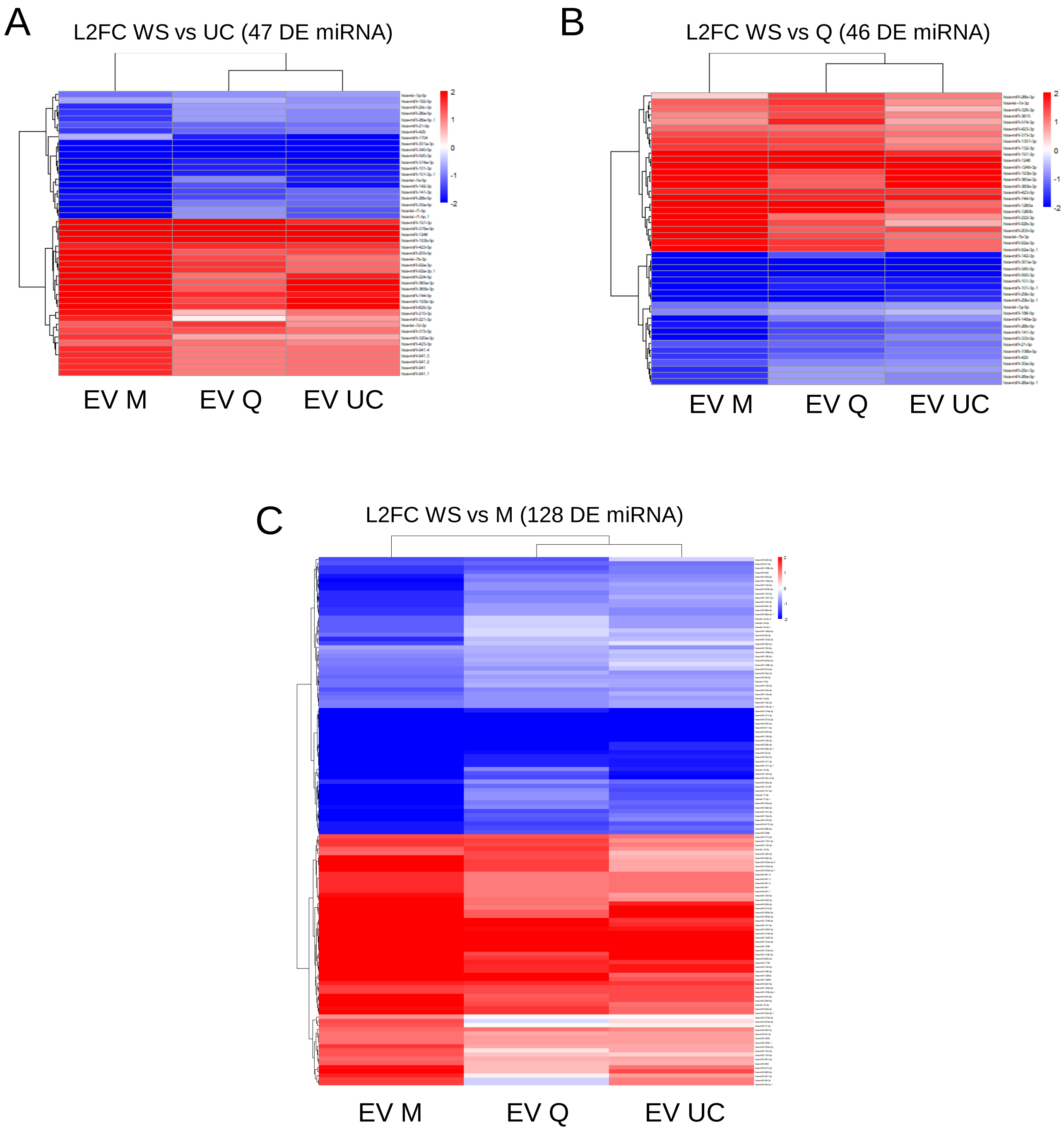

**Figure S4. Heatmaps of differentially expressed miRNAs between whole saliva and EV isolation methods.** Heatmaps showing the **$\log_2$ fold change ($Log_2FC$)** of significantly differentially expressed miRNAs (adjusted p-value < 0.05) in the following comparisons: (A) WS vs UC (47 miRNAs), (B) WS vs Q (46 miRNAs),(C) WS vs M (128 miRNAs).Each heatmap represents $Log_2FC$ values normalized on a scale from –2 to +2. The x-axis indicates the comparison between conditions; the y-axis lists differentially expressed miRNAs. Red indicates positive $Log_2FC$ values (upregulated in WS, downregulated in EVs), while blue indicates negative $Log_2FC$ values (upregulated in EVs, downregulated in WS). Heatmaps were generated using the pheatmap R package (version 1.0.12).

### Figure S5.

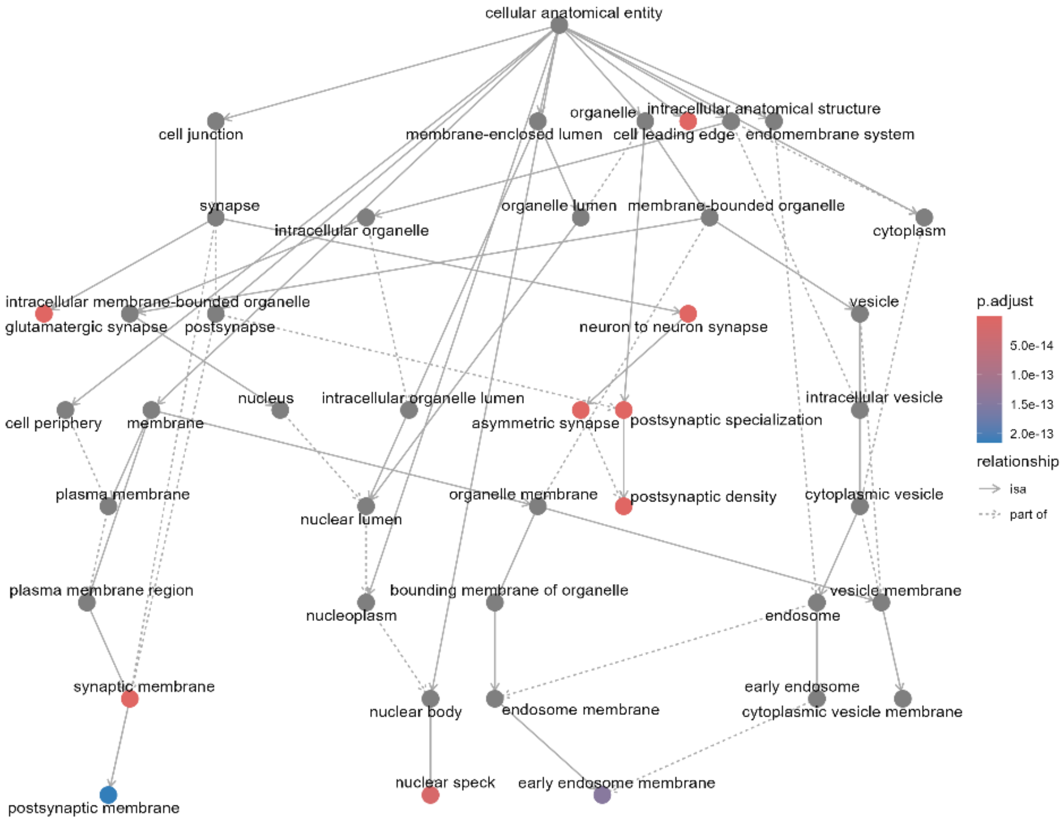

**Figure S5. Functional enrichment network of predicted gene targets in the Gene Ontology Cellular Component category.** Network graph showing Gene Ontology (GO) Cellular Component (CC) enrichment based on 14,635 predicted target genes of the 28 common and differentially expressed miRNAs identified in WS vs M, WS vs Q, and WS vs UC comparisons. Nodes represent enriched GO terms; solid arrows indicate hierarchical subcategories, and dashed arrows show broader category relationships. Red nodes: GO terms with low adjusted p-values (high significance). Blue nodes: GO terms with higher adjusted p-values (lower significance). Grey nodes: non-significant terms. The analysis was performed using the clusterProfiler R package (version 4.10.1).

Figure S6.

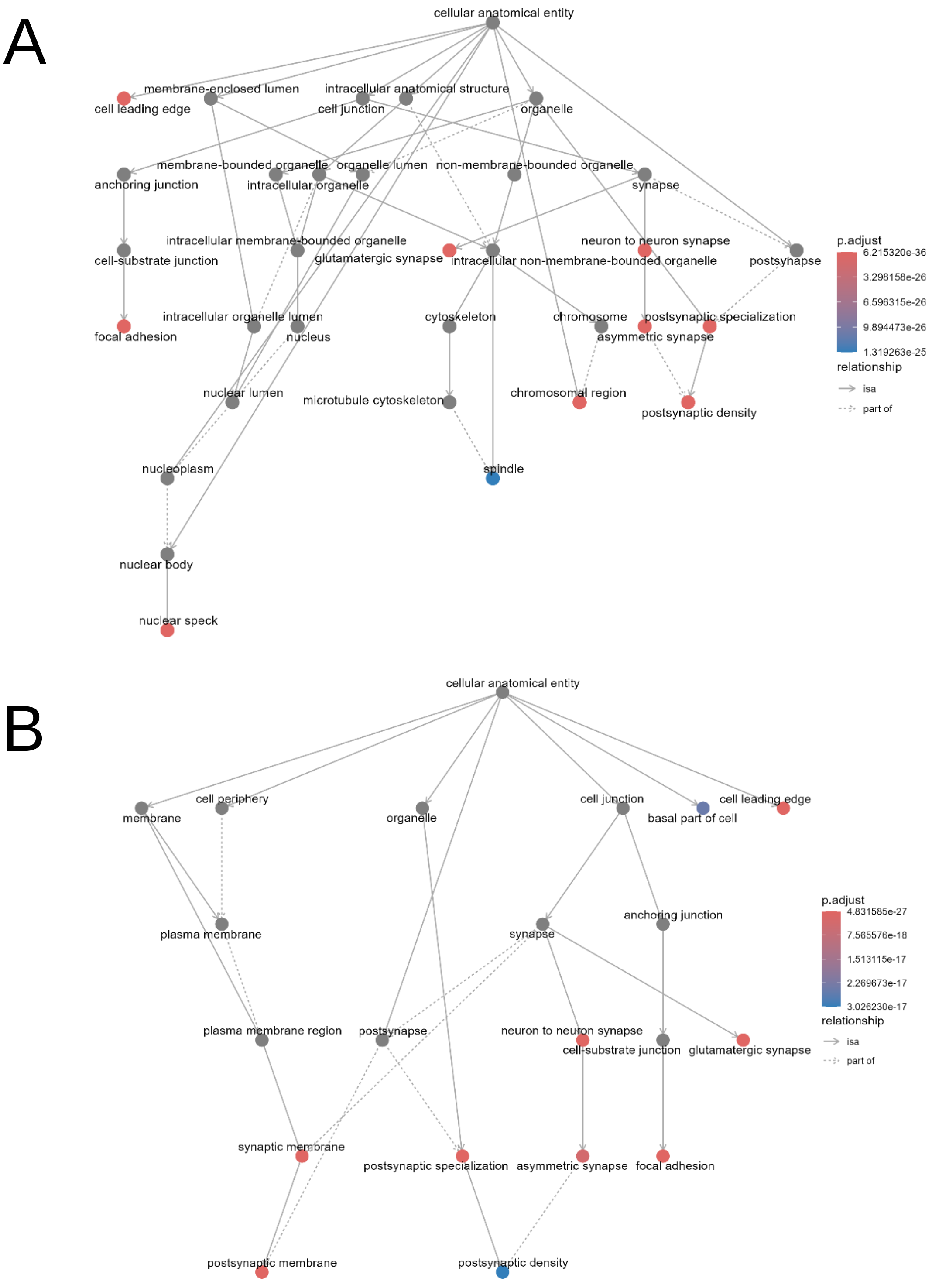

**Figure S6. Functional enrichment of gene targets of miRNAs differentially expressed in WS vs M.** (A) Network graph of Gene Ontology (GO) Cellular Component (CC) terms enriched among 16,397 validated target genes of the 65 miRNAs significantly differentially expressed between WS and M. (B) Network graph of GO-CC terms enriched among 17,674 predicted target genes of the same 65 miRNAs. Nodes represent enriched GO categories. Solid arrows indicate hierarchical subcategories. Dashed arrows show relationships between parent categories. Red nodes: highly significant GO terms (low adjusted p-values). Blue nodes: less significant GO terms (higher adjusted p-values). Grey nodes: non-significant terms. Functional enrichment analyses were performed using the clusterProfiler R package (version 4.10.1).

### Figure S7.

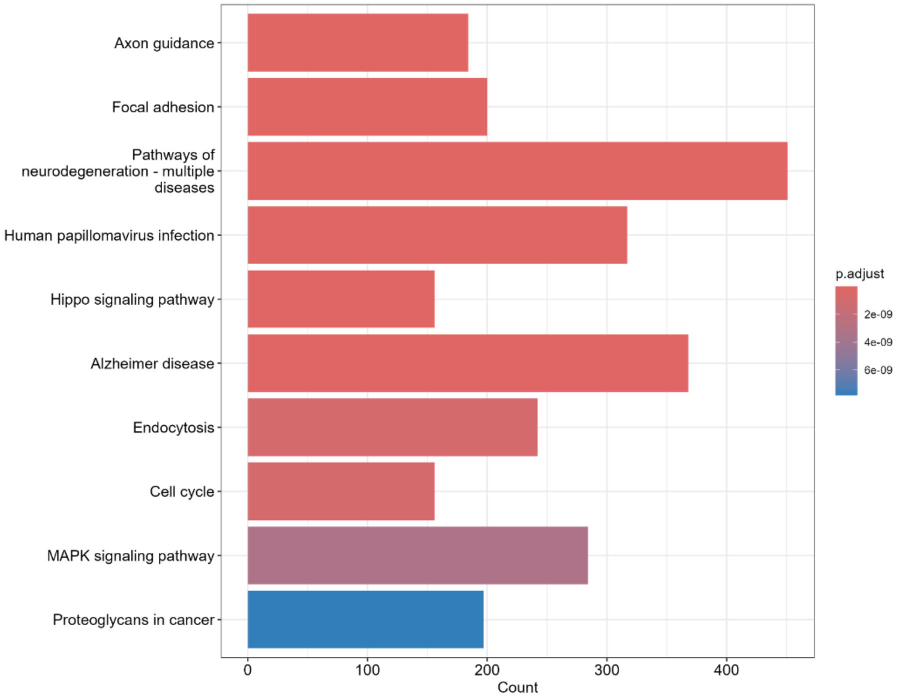

**Figure S7. KEGG pathway enrichment of validated target genes of M-specific miRNAs.** Bar chart showing KEGG pathway enrichment for 16,397 validated target genes of the 65 miRNAs uniquely differentially expressed in the M (immunoaffinity) condition. The x-axis indicates the number of genes associated with each pathway, while the y-axis lists the enriched KEGG pathways. The color gradient represents adjusted p-values: red indicates higher significance (low p-adjusted value), and blue indicates lower significance. Functional enrichment analysis was performed using the clusterProfiler R package (version 4.10.1) with the KEGG database (https://www.genome.jp/kegg/).

### Table S1 : List of miRNA used in the study

| Supplementary Table 1: List of miRCURY LNA miRNA PCR Assay | | |
|---|---|---|
| Name | Sequence | GeneGlobe ID |
| hsa-miR-423-5p | 5'UGAGGGGCAGAGAGCGAGACUUU | YP00205624 |
| hsa-miR-193-3p | 5'AACUGGCCUACAAAGUCCCAGU | YP00204591 |
| hsa-miR-26b-5p | 5'UUCAAGUAAUUCAGGAUAGGU | YP00204172 |
| hsa-let-7a-5p | 5'UGAGGUAGUAGGUUGUAUAGUU | YP00205727 |
